# Oriented Display of Cello-Oligosaccharides for Pull-down Binding Assays to Distinguish Binding Preferences of Glycan Binding Proteins

**DOI:** 10.1101/2023.01.26.525732

**Authors:** Markus Hackl, Zachary Power, Shishir P. S. Chundawat

## Abstract

The production of biofuels from lignocellulosic biomass using carbohydrate-active enzymes like cellulases is key to sustainable energy production. Understanding the adsorption mechanism of cellulases and associated binding domain proteins down to the molecular level details will help in the rational design of improved cellulases. In nature, carbohydrate-binding modules (CBMs) from families 17 and 28 often appear in tandem appended to the C-terminus of several endocellulases. Both CBMs are known to bind to the amorphous regions of cellulose non-competitively and show similar binding affinity towards soluble cello-oligosaccharides. Based on the available crystal structures, these CBMs may display a uni-directional binding preference towards cello-oligosaccharides (based on how the oligosaccharide was bound within the CBM binding cleft). However, molecular dynamics (MD) simulations have indicated no such clear preference. Considering that most soluble oligosaccharides are not always an ideal substrate surrogate to study the binding of CBMs to the native cell wall or cell surface displayed glycans, it is critical to use alternative reagents or substrates. To experimentally assess any binding directionality of CBMs towards soluble cello-oligosaccharides, we have developed a simple solid-state depletion or pull-down binding assay. Here, we specifically orient azido-labeled carbohydrates from the reducing end to alkyne-labeled micron-sized bead surfaces, using click chemistry, to mimic insoluble cell wall surface-displayed glycans. Our results reveal that both family 17 and 28 CBMs displayed a similar binding affinity towards cellohexaose-modified beads, but not cellopentaose-modified beads, which helps rationalize previously reported crystal structure and MD data. This indicates a preferred uni-directional binding of specific CBMs and could explain their co-evolution as tandem constructs appended to endocellulases to increase amorphous cellulose substrate targeting efficiency. Overall, our proposed workflow can be easily translated to measure the affinity of glycan-binding proteins to click-chemistry based immobilized surface-displayed carbohydrates or antigens.

## 1 Introduction

The cost-effective breakdown of lignocellulose biomass waste to fermentable sugars and its subsequent fermentation to ethanol is a crucial process for the production of sustainable fuel in the future (1). In this process, enzymatic hydrolysis of lignocellulosic biomass is an important step and relies on the effective deployment of a mixture of enzymes to hydrolyze the biomass-derived polysaccharides into fermentable sugars (2). Many of those carbohydrate-active enzymes (CAZymes) are multi-domain polypeptides where a single or multiple CBMs are attached to one or more catalytic domains (CDs). The CBM is responsible for the recognition of and binding to the substrate, whereas the CD breaks down the substrate into shorter oligosaccharides or fermentable monosaccharides for direct cellular uptake (3). The CBM, therefore, plays a pivotal role in the depolymerization process since it is often the main driver for substrate recognition and targeting specific regions of the cell wall polysaccharides like cellulose (4, 5). Based on specific substrate affinity, CBMs can be categorized into three groups. Type A CBMs bind to crystalline regions of cellulose, whereas type B and type C CBMs bind to oligosaccharide chains or single monosaccharide units, respectively (6).

The binding sites of type B CBMs range from a deep binding groove, as seen in CBM4 (7–10), to a relatively shallow groove as seen in CBM families 17 and 28 (11–13). Type-B CBMs like CBM 17 and 28 can accommodate between 3 and 6 glucopyranose units of a cello-oligosaccharide within the binding cleft. While type B CBMs exhibit a stronger affinity for insoluble amorphous cellulose compared to Avicel® or microcrystalline cellulose (9, 14–17), these CBMs are also reported to bind short cello-oligosaccharides (9, 11, 14, 16, 18–20). However, the CBM-glycan binding affinity drops with decreasing chain length of the cello-oligosaccharide. Crystal structures of CBMs from families 17 and 28 containing a bound ligand show that the cello-oligosaccharide is actually bound in the opposite direction for each as shown in Figure 1, although the binding free energies as estimated experimentally were similar for both protein families for the same cello-oligosaccharide (11–13). MD simulations have been carried out to investigate whether there is any preference for the direction of the ligand docked in the type-B CBM binding pocket (10, 21, 22). The MD study for CBM 17 and 28 (21) revealed that while the cellopentaose ligand was in contact with the CBM over the entire MD simulation period for any orientation of the ligand, not all orientations exhibited equally well-stabilized protein-ligand interactions. In fact, the root mean square fluctuations (RMSF) were around 1 Å for CBM17 binding the ligand from the reducing end and CBM28 binding from the non-reducing end of cellopentaose. The opposite binding orientations (i.e. CBM17 binding from the non-reducing end) showed more than twice as much RMSF as well as sliding of the cellopentaose in the binding pocket, indicating less stable and potentially weaker binding interactions (21).

**Figure 1:**
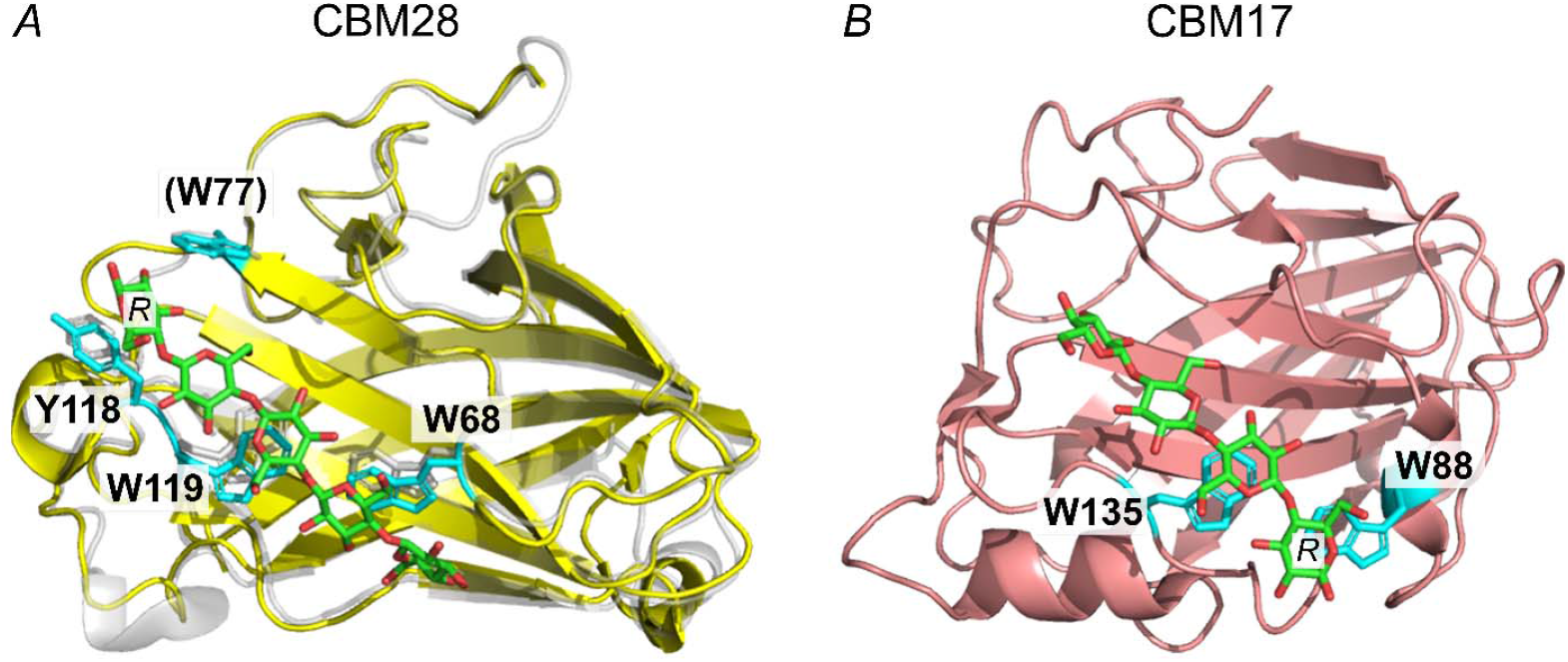
Orientation of cello-oligosaccharides in the binding pocket of type B CBMs. The reducing end of the oligo is labeled “R” and aromatic residues in close contact are highlighted in cyan. The cello oligomers are oriented in different directions in the binding pockets. A) No crystal structure of A. akabai CBM28 bound to a ligand exists, hence CBM28 bound to cellopentaose from R. josui (PDB ID 3aci, light grey) was aligned with A. akabai CBM28 (PDB ID 1uww, yellow). Based on structural similarities, potential residues of A. akabai CBM28 are noted. W77 may not be involved in ligand binding with this alignment but may aid in an alternative binding orientation (13). B) CBM17 from C. cellulovarans bound to cellotetraose (PDB ID 1j84).

Bacterial cellulases such as Cel5A from *Alkalihalobacillus akibai* (formerly known as *Bacillus sp*. 1139) (18), Cel5A from *Ruminoclostridium josui* (14), and Cel9B from *Cellulomonas fimi* (23) contain type B CBMs in tandem. *A. akabai* and R. *josui* native tandem design consists of CBM17/CBM28, whereas *C. fimi* tandem design is constructed of two CBM4. It is hypothesized, that multiple type B CBMs in tandem can help bind different regions of the insoluble and highly disordered cell wall substrate (15). Amorphous regions of cellulose are characterized by a reduced crystallinity and degree of polymerization, although structural details remain obscured (24). It was shown that different type B CBMs bind to different regions of cellulose and cell walls non-competitively, indicative of recognizing different binding sites on the substrate (25–27). Though, the identification of those different binding sites is difficult to achieve due to the complex nature of the insoluble substrate. Using well-defined substrates, such as cello-oligosaccharides, the binding affinity of CBMs can be accurately determined with isothermal titration calorimetry (ITC), fluorescence or ultraviolet (UV) absorption (11, 14, 16, 18–20). However, it is not possible to infer information about the binding configuration of the soluble oligosaccharide ligand in the CBM binding pocket, and the only structural information about ligand orientation stems indirectly from examining crystal structures of CBMs with a bound ligand.

To address the open question about experimental verification of cello-oligosaccharide ligand binding orientation in type B CBMs, we developed a well-defined ‘amorphous’ cellulosic substrate surrogate to perform solid-state depletion assays. Micron-sized polystyrene (PS) beads were functionalized with cello-oligosaccharides that were oriented with a defined stereochemistry (i.e., with the non-reducing end exposed and available for interacting with the solvent). To our knowledge, this is the first solid-state depletion assay using soluble cello-oligosaccharides with a degree of polymerization (dp) >4. An overview of the PS-bead preparation scheme is outlined in Figure 2. Briefly, we use Shoda’s reagent (28) to convert cello-oligosaccharides into corresponding glycosyl azides (Figure 2-A). Micron-sized PS beads were functionalized with dibenzocyclooctyne (DBCO) (Figure 2-B) and the glycosyl azides were covalently linked to these beads using click-chemistry, creating a proxy insoluble cellulosic substrate with defined properties.

**Figure 2:**
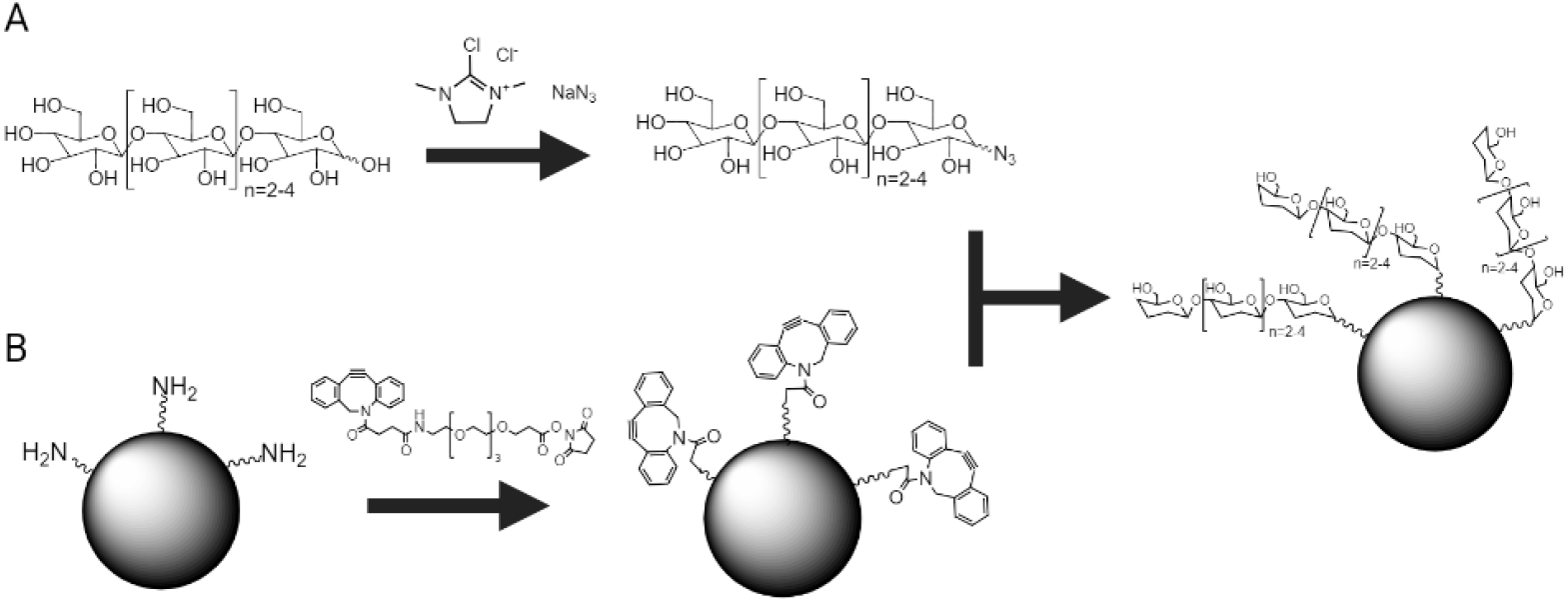
Overview of the method to immobilize soluble cello-oligosaccharides to PS beads using click chemistry to create defined solid substrates. A) The conversion of cello-oligosaccharides to corresponding glycosyl azides using Shoda’s reagent (DMC or 2-chloro-1,3-dimethylimidazolinium chloride) and sodium azide. B) Conversion of amino groups displayed on micron-sized PS beads to DBCO groups using a PEGylated NHS-ester. The glycosyl azides are then reacted with the DBCO to create cello-oligosaccharide-modified PS beads as the final product.

The cello-oligosaccharide functionalized PS beads were then used in a solid-state depletion assay to determine the binding properties of CBM28 from *A akabai* and CBM17 from *C. cellulovarans. Both* CBMs were fused to a green fluorescent protein (GFP) tag for protein quantification. Our results show that there is indeed a difference in the equilibrium binding affinity of CBM17 and CBM28 towards cellopentaose-modified PS beads, indicating a preferred binding orientation for each protein. However, no significant difference was found for cellohexaose-modified beads. Our method can be applied to immobilize any soluble carbohydrates or oligosaccharides via the reducing end, thus creating an insoluble substrate analog to investigate protein binding at interfaces through easy-to-execute pull-down assays. More advanced characterization methods such as quartz crystal microbalance (QCM) or surface plasmon resonance (SPR) can utilize the same workflow, once the QCM/SPR surface is functionalized with DBCO moieties, thus highlighting the versatility of the described workflow.

## 2 Material and Methods

Unless otherwise mentioned, all reagents were either purchased from VWR International, USA, Fisher Scientific, USA, or Sigma-Aldrich, USA. Cello-oligosaccharides were either purchased from Biosynth Carbosynth, USA or Neogen, USA. Amino-modified polystyrene beads (AP-30-10) with a nominal diameter of 3.4 µm were purchased from Spherotech Inc, USA.

### 2.1 Protein expression and purification

The genes for *Clostridium cellulovarans* CBM17 and *Alkalihalobacillus akabai* CBM28, codon optimized for *E*.*coli*, were obtained from Genewiz, USA, and expressed and purified as GFP-fusion constructs (see Supplemental Figure S1) in *E. coli* BL21-CodonPlus(DE3)-RIPL (Stratagene, USA) as described previously (29). The sequences are summarized in Supplemental Table S1.

### 2.2 Azide modification of cello-oligosaccharides

The anomeric hydroxy group of the cello-oligosaccharide was substituted with an azide group following the steps outlined by Tanaka et al (30). The reaction mixture composition per 10 mg of the substrate is summarized in Table 1. First, the cello-oligosaccharide was dissolved in the respective amount of heavy water (D_2_O) and transferred to a 20 ml screw-capped glass vial containing a small magnetic stir bar. Next, sodium azide (NaN_3_) and 2-chloro-1,3-dimethylimidazolinium chloride (DMC) were added, followed by the addition of N,N-diisopropylethylamine (DIPEA). The reaction mixture was stirred for 30 minutes at room temperature and quenched with twice the reaction mixture volume of deionized (DI) water. Immediately after quenching, the mixture was transferred to a dialysis membrane (Spectrum™ 131060, MWCO 100-500 Da) and dialyzed at room temperature against DI water for 90-170 hours (4-7 days) with replacement of water every 8-24 hours (31). To prevent microbial growth, ProClin™ 300 at 0.05% (v/v) was added between hours 36-84. The final dialysis step (last 8-12 hours) did not contain any additives. After dialysis, the liquid was transferred to a 50 ml conical flask and the water evaporated *in vacuo*. Finally, the solid residue was dissolved in 1 ml of DI water.

**Table 1:** Reaction mixture composition per 10 mg of cello-oligosaccharide to generate glycosyl azides

|  | D-Cellotetraose | D-Cellopentaose | D-Cellohexaose |
| --- | --- | --- | --- |
| D<sub>2</sub>O (ml) | 0.5 | 1.0 | 2.0 |
| NaN<sub>3</sub> (mg) | 81.3 | 162.6 | 325.2 |
| DMC (mg) | 10.6 | 21.2 | 42.4 |
| DIPEA (ml) | 0.065 | 0.066 | 0.132 |

The conversion and overall yield were quantified through densitometric analysis of thin-layer chromatography (TLC) images. Aluminum-backed TLC silica gel 60 F_254_ plates (Supelco^®^ 1.05554.0001) were spotted with the dialyzed, resuspended reaction mixture and unmodified cello-oligosaccharide control and developed using a mobile solvent mixture of butanol-ethanol-water in a volumetric ratio of 5-5-4 (32). After TLC, the dried plates were sprayed with 0.1% orcinol (in 1.8 M sulfuric acid in 190 proof ethanol), dried, and developed at 100°C for 3-5 minutes until the cello-oligosaccharide spots turned dark.

The conversion was calculated as 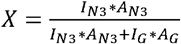 where is the pixel intensity of the *I*_*N*3_ azide-modified cello-oligosaccharide (glycosyl azide), measured over the spot size *A*_*N*3_ and *A*_*G*_ represent the mean intensity and spot size of the unmodified cello-oligosaccharide. The concentration was determined by creating a standard curve of the unmodified cello-oligosaccharide at concentrations between 1 mM to 0.1 mM and comparing the intensity of the reaction mixture to the standard curve. The overall yield was calculated as 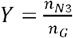, where *n*_N3_ is the final amount of glycosyl azide and *n*_*G*_ is the initial amount of substrate (unmodified cello-oligosaccharide) added.

### 2.3 Cello-oligosaccharide functionalization of amino-modified PS beads

The steps outlined here describe the preparation of one sample (one data point) for the solid-state depletion assay and scale-up depends on the number of samples required. The control samples for non-specific binding receive the same treatment, except during the click-reaction step, phosphate buffered saline (PBS) was used instead of glycosyl azides. It is important to ensure that all buffers during the functionalization steps are free of azides as it competes with the click reaction of glycosyl azides. The glycosyl azides are covalently linked to the amino-functionalized beads in a two-step process as shown in Figure 2. In the first step, the amino groups on the PS bead surface were converted into DBCO groups. First, 20 µl of bead stock solution (concentration of NH_2_ groups is approximately 250 µM as per the manufacturer’s specification) were spun down and resuspended in PBS at pH 7.4. To functionalize the beads with DBCO, a PEGylated NHS-ester linker was used (i.e., DBCO-polyethylene glycol (PEG(4)) N-hydroxysuccinimidyl (NHS) ester). The beads were resuspended in 20 µl of 250 µM DBCO-PEG(4)-NHS ester (linker) in PBS and incubated on a rotisserie overnight at room temperature. The conversion to DBCO moieties was confirmed by analyzing the fluorescence intensity of single beads functionalized with azide-labeled fluorophores (see Supplemental Figures S2 and S3). The beads were washed three times in 100 µl PBS to remove any unreacted linker. In the second step, the glycosyl azides were covalently attached to the DBCO moieties displayed on the PS beads. The glycosyl azide solution was diluted to 0.5-1 mM in PBS by adding the respective volume of water and 10x PBS concentrate. Next, the DBCO-modified beads were resuspended in 20 µl of glycosyl azide-containing buffer and incubated on a rotisserie overnight. Finally, the functionalized beads were washed three times in PBS and used the same day for the solid-state depletion assay.

### 2.4 Solid-state depletion or pull-down binding assay

The solid-state depletion assay follows the general steps used often to characterize CBM binding to insoluble substrates like microcrystalline or amorphous cellulose (33, 34). The working buffer (WB) used in the binding experiments was 10 mM PBS at pH 7.4 containing 0.2 mg/ml of bovine serum albumin (BSA) and Pluronic-F127, respectively. The CBMs were diluted in WB to a concentration range between 50 and 1000 nM. First, the bead samples (20 µl each) were resuspended in PCR tubes containing 100 µl of WB and incubated on a rotisserie for 15 minutes to passivate the bead surface. After centrifugation (2000 x g for 1-3 minutes) and removal of the supernatant, the beads (cello-oligosaccharide-functionalized and non-specific binding control) were resuspended in 100 µl of the prepared CBM dilutions and incubated on the rotisserie for 120 minutes at room temperature. Next, the beads were centrifuged and 90 µl of supernatant was transferred to a new PCR tube, which was spun down again. Finally, 80µl of this supernatant was transferred to a black, clear bottom 96-well plate for unbound protein fluorescence quantitation. Two separate centrifugation steps were necessary to reduce the interference from beads being accidentally transferred to the 96-well plate.

The CBM concentration was determined by measuring the fluorescence signal of the appended GFP domain. The GFP-CBM standard curve was prepared from the same CBM dilutions used in the solid-state depletion assay and the fluorescence was quantified in a spectrophotometer (SpectraMax M5e, Molecular Devices) using 480 nm excitation, 512 nm emission, and a cut-off of 495 nm.

To obtain the free protein concentration, the readings of the non-specific binding samples at the same protein concentration were averaged. The bound protein concentration was determined by subtracting the free protein concentration from each cello-oligosaccharide-modified bead reading. Instead of using the mass of substrate added, the amount of bound protein was based on the number of theoretically available binding sites on the beads. Assuming a 100% conversion during both steps of the bead functionalization, this results in 5 nmol of total available binding sites per 20 µl of beads.

The Langmuir one-site binding model was used to determine the dissociation constant, *K*_d_, and available binding sites on the cello-oligosaccharides functionalized bead surface, *n*_*max*_. The model equation can be written as 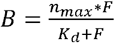, where B represents the concentration of bound protein and F the concentration of free/unbound protein. The model parameters were estimated using OriginPro 2020® using the built-in Levenberg-Marquardt algorithm.

## 3 Results

### 3.1 Azide modification of cello-oligosaccharides

The successful conversion of cello-oligosaccharides (i.e., cellotetraose, cellopentaose, and cellohexaose) to corresponding glycosyl azides was verified using TLC and representative results are shown in Figure 3. While there is only a small separation between the cello-oligosaccharides in the standard (Lane 1), the glycosyl azides separated well from the unmodified cello-oligosaccharides (Lanes 2-4, the red arrow indicates glycosyl azide). Using dialysis, it is not possible to separate the unmodified substrate from the product due to only a minor difference in molecular weight. However, dialysis efficiently removes the free azides, which would significantly interfere with the subsequent click reaction (see Supplemental Figure S4). Unmodified cello-oligosaccharides will be removed at washing steps after the click reaction, hence no separation between starting unreacted substrate and glycosyl azide was necessary.

**Figure 3:**
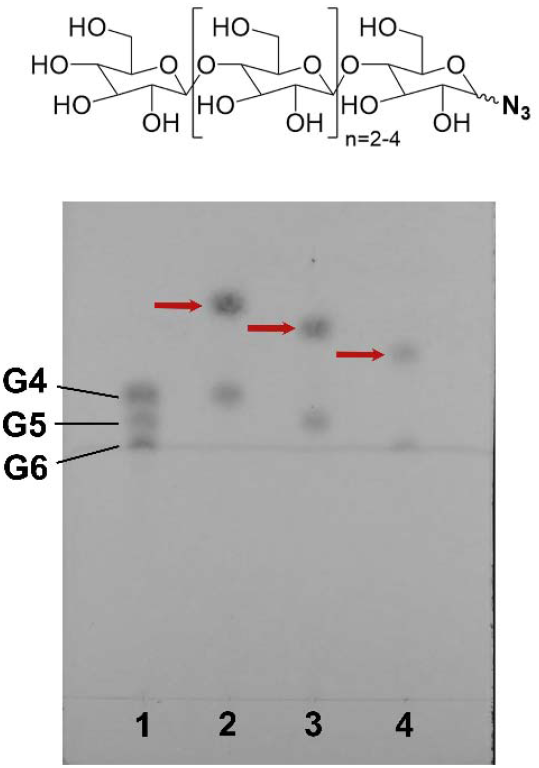
Thin-layer chromatography confirms the conversion of cello-oligosaccharides to azide-modified cello-oligosaccharides. The glycosyl azide products formed are indicated by the red arrow. The common structure of a glycosyl azide is shown on the top. Lane 1: Standard ladder of cellotetraose (G4), cellopentaose (G5), and cellohexaose (G6). Lane 2: Cellotetraose and cellotetraosyl azide (n=2). Lane 3: Cellopentaose and cellopentaosyl azide (n=3). Lane 4: Cellohexaose and cellohexaosyl azide (n=4).

The conversion was 84% for cellotetraose, while cellopentaose and cellohexaose showed a conversion of 59% and 63%, respectively. The total product yield from the reaction mixture after purification increased with the molecular weight of the carbohydrate and is 2, 6, and 9% for cellotetraose, cellopentaose, and cellohexaose respectively. The yield is significantly lower than previously reported (30), most likely due to sample loss during dialysis instead of using High-Performance Liquid Chromatography (HPLC) as the purification method. Even though the cutoff of the membrane is less than 500 Da, most of the product is lost during the dialysis because it either diffuses through the pores and/or irreversibly binds to the cellulose ester membrane. Nevertheless, the removal of free azides using dialysis is a viable alternative in case HPLC at the preparative scale is not readily accessible.

### 3.2 Solid-state depletion protein-ligand binding assay

Image analysis of single beads revealed that CBMs bind specifically (p<0.05, n=50) to cello-oligosaccharide functionalized beads as shown in Supplemental Figure S5. The binding data and fitted Langmuir one-site binding model are summarized in Figure 4 and the region of less than 100nM of the binding isotherms is shown in Supplemental Figure S6. The dissociation constants and the maximum number of available binding sites are reported in Table 2. The p-values for each parameter estimation are summarized in Supplemental Table S2. The dissociation constants as determined by our assay method were 40x-2200x lower compared to the results reported for the same proteins and substrates using ITC (11, 18). The main difference is that the substrate or ligand can freely diffuse in solution during ITC and is not immobilized to a solid surface. A reduction of the dissociation constant (i.e., or equivalent increase in binding affinity) has been previously reported for immobilized antibodies (35) towards respective antigen ligands. A similar phenomenon could explain the higher binding affinity seen for CBMs towards surface-immobilized oligosaccharides in our pull-down assay versus ITC assays using soluble ligands.

**Table 2:**
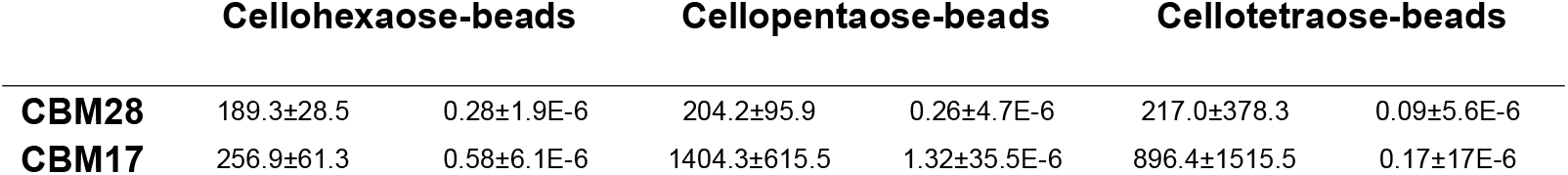
Summary of Langmuir one site binding model fit parameters for CBM17 and CBM28 on cello-oligosaccharide-modified PS beads. The values indicate mean ± SE. The unit of is (nM), the unit of is nmol protein/nmol NH_2-eq_.

|  | Cellohexaose-beads |  | Cellopentaose-beads |  | Cellotetraose-beads |  |
| --- | --- | --- | --- | --- | --- | --- |
| **CBM28** | 189.3±28.5 | 0.28±1.9E-6 | 204.2±95.9 | 0.26±4.7E-6 | 217.0±378.3 | 0.09±5.6E-6 |
| **CBM17** | 256.9±61.3 | 0.58±6.1E-6 | 1404.3±615.5 | 1.32±35.5E-6 | 896.4±1515.5 | 0.17±17E-6 |

**Figure 4:**
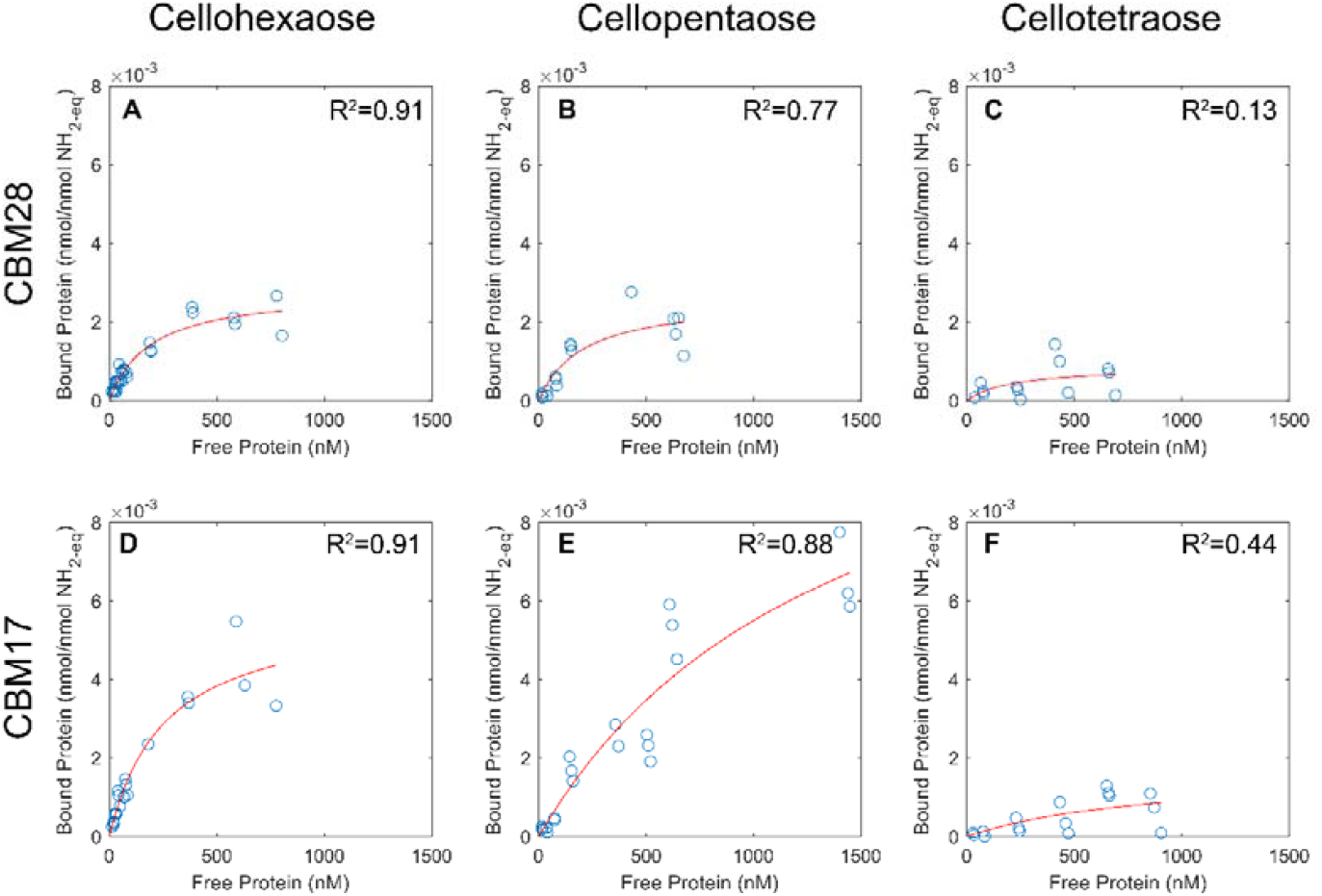
Binding data (blue circles) and fitted isotherms (red line) of CBM28 (top) and CBM17 (bottom) for cello-oligosaccharide-modified beads. The R^2^-value of the isotherm fit is shown in the top right corner of each panel. While the binding to cellohexaose-modified beads is similar for CBM17 and CBM28, there is a clear difference in binding for cellopentaose-modified beads. Both proteins show marginal binding to cellotetraose-modified beads. Panel A-C) binding data for CBM28 o cellohexaose, cellopentaose, and cellotetraose-modified beads respectively. Panel D-F) binding data for CBM17 on cellohexaose, cellopentaose, and cellotetraose-modified beads respectively.

Both CBMs display a similar dissociation constant for cellohexaose-modified beads, however, the maximum number of binding sites is twice as high for CBM17. Differences in binding affinity become more evident when comparing the results for cellopentaose-modified beads. While the dissociation constant for CBM28 on cellopentaose-modified beads is similar to cellohexaose-modified beads, CBM17 displays a ∼5.5-fold increase in the dissociation constant but a ∼2.3-fold increase in the number of binding sites. Although the CBM17 concentration was extended to 1.5 µM, the binding curve of CBM17 on cellopentaose-modified beads did not fully level off as it was the case for CBM28. This, along with the increase in dissociation constants, indicates that CBM17 displays weaker binding towards cellopentaose-modified beads in our assay.

## 4. Discussion

We report the development of a solid-state depletion assay for testing protein binding to PS bead surface-immobilized glycans as an alternative method to characterizing protein binding to soluble glycans. The modular approach using amino-modified surfaces will enable the adoption of the azido functionalization process to other analytical techniques such as QCM or SPR. The key step is the conversion of soluble glycans to glycosyl azides using Shoda’s reagent, which can be adopted for various carbohydrates, such as xylo-or malto-oligosaccharides and even complex, branched sugars (30). In particular, the directed immobilization of complex carbohydrates as found in mammalian systems may be of great interest. Most of those glycans are immobilized on either antibodies or cell surfaces, which restrict their conformation and facilitate specific antibody-antigen interactions (36). Alternative chemical methods to site-specifically functionalize carbohydrates for immobilization on surfaces are time-consuming and involve tedious protection/deprotection steps of hydroxy groups (37, 38), thus the use of Shoda’s reagent is a promising and simple alternative. Amino-modified beads are commercially available and other QCM/SPR sensor surfaces made of quartz or borosilicate glass can be amino-functionalized with aminated silanes (39–41). The functionalization of amino-modified surfaces with DBCO moieties is straightforward and can easily be verified using azide-modified fluorophores as shown in Supplemental Figure S3. However, the removal of free azides from Shoda’s reaction mixture is critical, as even a 0.1x molar excess of free unreacted azide can reduce click-chemistry reaction efficiency by more than 30% (see Supplemental Figure S4).

Previous MD simulations have revealed that the root-mean-square fluctuations or RMSF of cellopentaose upon binding to CBM17 and CBM28 can vary significantly depending on the ligand binding orientation (21). Based on those simulations, CBM17 seems to prefer binding from the reducing while CBM28 prefers binding from the non-reducing end. The directed immobilization of cello-oligosaccharides in our bead assay displays the non-reducing end for binding. This could limit protein access to favorable (more stable) ligand binding sites for CBM17, although the total number of available binding sites increases significantly compared to CBM28. In contrast, CBM28 prefers binding to oligosaccharides from the non-reducing end, and as such only exhibits favorable binding interactions with the immobilized ligands, which was not detected using soluble substrates previously. In fact, the dissociation constant and number of binding sites for CBM28 are similar for cellohexaose- and cellopentaose-modified beads. This is the first reported evidence showcasing the directional cello-oligosaccharide binding preference of CBM 17 vs. 28 families.

Both CBMs showed poor binding towards to cellotetraose-modified beads as seen in Figure 4, panels C and F as well as in the large error in the binding parameters data reported in Table 2 and p-values in Supplementary Table S2. This may be because the covalent linkage through the DBCO moiety could impose a steric hindrance that effectively reduces the total number of pyranose rings available to engage via suitable hydrogen bonding and stacking interactions with the residues in the protein binding cleft. In other words, the cellohexaose-modified bead could be an effective cellopentaose-modified bead, and so on. This hypothesis may be supported by the fact that both CBM17 and CBM28 do not bind or just weakly bind to cello-oligosaccharides of a dp <4 (14, 16, 18). However, *A. akabai* CBM28 has a surface-exposed tryptophan at position 77 (see Figure 1, which may aid in binding cello-oligosaccharides in an alternative binding mode (12). In addition, CBM17 also lacks one aromatic residue in the binding pocket when compared to CBM28, which could explain the reduction in binding affinity as determined in our assay.

Previous work has shown that strict control of antibody orientation resulted in 100-fold stronger antigen−antibody complexation than controls (35). Analogously, it is likely that a reduction in the total degrees of freedom available for the surface-displayed ligands and well-defined ligand orientations could also result in tighter protein-ligand binding interactions at the solid-liquid interface. Similarly, type-B CBMs have been reported to show 10-to-100-fold higher binding affinity towards insoluble amorphous cellulose versus soluble cello-oligosaccharides (42). It was hypothesized previously that the high-affinity binding site interactions of CBMs with insoluble versus soluble ligands are due to relative gains in binding enthalpy (ΔH) and not gains seen in configurational entropy (ΔS). In contrast with results reported previously for CBM17/28 binding at high-affinity sites, the low-affinity binding interactions did show a gain in configurational entropy with a compensating enthalpic loss. However, since the structure of insoluble crystalline or amorphous cellulose is unknown, it has been challenging in the past to directly associate energetic observations made from ITC or pull-down binding assays with structural features of the ligand. Using well-defined oligosaccharides immobilized to PS-beads, it would be now possible to systematically explore the binding interactions of proteins with surface-immobilized glycans.

The surface density of amino groups on the beads is approximately 1.7 NH_2_/Å^2^. Assuming that all amino groups are converted to DBCO groups for click-chemistry labeling to cello-oligosaccharides and that one CBM covers around 50 Å^2^, then around 165 cello-oligosaccharides would be close to one CBM. Our binding data suggests that less than 1% of the available NH_2_ groups result in a successful CBM binding event. Steric hindrance from the PEG(4)-DBCO linker as well as incomplete reactions could significantly reduce the total number of cello-oligosaccharides close to each binding site. Nevertheless, it may be possible that *A. akabai* CBM28 may be able to engage with more than one cello-oligosaccharide at once due to W77, which is absent in CBM17 as well as *R. josui* CBM28. Future experiments could be carried out in which W77 is substituted to alanine or glycine and/or the binding site is further modified by replacing Y118 with alanine as well to only leave two aromatic residues in the binding pocket of CBM28. Nevertheless, our assay revealed a significant difference in dissociation constants if the substrate is displayed in an oriented manner. This could shed light on the reason for CBM17 and CBM28 to naturally occur in tandem.

## Acknowledgments

S.P.S.C. acknowledges support from the NSF (CBET CAREER Award 1846797) and Rutgers SOE Startup Funds. Partial support was provided by NSF CBET Award 1704679.

