## Supplementary Information for "Oriented Display of Cello-Oligosaccharides for Pull-down Binding Assays to Distinguish Binding Preferences of Glycan Binding Proteins" (from another version of this preprint: version 2, 2023-02-08)

**This PDF file includes**:

Supplementary text
Supplementary Figures S1 to S6
Supplementary Tables S1 to S2
SI References

### Gene sequences for CBM17 and CBM28

As shown in Supplemental Figure S1, both CBMs were fused to $His_8$-GFP separated by a flexible linker domain as previously described (1). The gene sequences for linker and CBM can be found in Supplemental Table S1.

### Single polystyrene (PS) bead analysis using wide-field fluorescence microscopy

To verify the proposed reactions and binding events, a method to analyze single PS beads using fluorescence microscopy was developed. Fluorescence images were taken at 20x magnification with a scientific camera (Kiralux, Thorlabs Inc.) using µManager (2) on an inverted fluorescence microscope (Olympus IX 71) equipped with the necessary filters to monitor green fluorescence. The individual steps in image processing and analysis were carried out in Fiji (3) and are shown in Supplemental Figure S2. Each image was first corrected for background and shading (4), followed by thresholding to remove the background. The exact values for thresholding depend on the signal/noise ratio of the particular experiment and were set such that the outline of single beads was clearly visible. Finally, 50 single beads were randomly chosen that were not in close proximity to other beads and the mean value was recorded.

### Confirmation of DBCO linker functionalization of aminated PS beads

The functionalization of aminated beads with the NHS-PEG(4)-DBCO linker was confirmed by using an azide labeled fluorophore ($N_3$-488, or Rhodamine 110-azide) which reacts with the DBCO moiety on the beads. A schematic overview of the experiment is shown in Supplemental Figure S3-A. Image analysis was carried out for single beads as described above. It is important that all buffers are free of amines or azides as that would interfere with the functionalization steps. First, 20 µl of aminated beads were washed three times in 100 µl PBS pH 7.4 and equally split into two PCR tubes. Next, one sample of beads was resuspended in 10 µl PBS containing 250 µM NHS-PEG(4)-DBCO, whereas the other one was resuspended in PBS only. The beads were incubated on a rotisserie overnight at room temperature, followed by three washes of 100 µl PBS each. Next, both samples were resuspended in 10 µl PBS containing 10 µM of $N_3$-488 and incubated on a rotisserie for 120 minutes. Finally, both samples were washed three times with 100 µl PBS and imaged on the fluorescence microscope. As it can be seen in Supplemental Figure S3-B, the addition of the NHS-PEG(4)-DBCO linker results in a significantly higher average pixel intensity compared to the control, thus verifying that the beads were successfully functionalized with DBCO moieties for subsequent click reactions.

### Influence of free azides on click reaction of DBCO-functionalized beads

The conversion of carbohydrates to glycosyl azides requires the presence of 50-120x molar excess of sodium azide. The removal of those free azides from the reaction mixture is important since the free azides compete with the click reaction of glycosyl azides. To assess the influence of free azides on the click reaction, the $N_3$-488 reaction described above was carried out in the presence of varying amounts of sodium azide. The functionalization of beads with DBCO and all washing steps and analysis were identically carried out as described above. The only difference was that $N_3$-488 was diluted to 10 µM in PBS containing 100x, 10x, 1x, 0.1x, and 0x molar excess of sodium azide with respect to $N_3$-488. As shown in Supplemental Figure S4, even a 0.1x molar excess of free sodium azide can significantly reduce the obtained fluorescence signal. The images were taken at 300 ms exposure time and no fluorescence signal was detected for the 100x sample at this setting.

**Binding specificity of green fluorescent protein (GFP)-CBM28 to cello-oligosaccharide functionalized beads**

Bead samples were functionalized with DBCO as described above. Instead of using $N_3$-488, the bead samples were resuspended in 10 µl PBS containing either 250 µM cellopentaosyl azide, 250 µM cellopentaose, or only PBS and allowed to mix on a rotisserie over-night at room temperature. Next, the beads were washed three times in 100 µl PBS and resuspended in 100 µl working buffer (WB, PBS containing 0.2 mg/ml of BSA and Pluronic-F127) to passivate the surface for 15 minutes. Following passivation, the beads were spun down and resuspended in 10 µl WB containing 1 µM of GFP-CBM28 and incubated for two hours at room temperature on a rotisserie. Finally, the beads were washed twice with 100 µl WB, resuspended in 10 µl WB and the samples were imaged and analyzed as described above. Supplemental Figure S5 summarizes the normalized mean pixel intensity from multiple beads. The obtained fluorescence from a bead sample functionalized with cellopentaosyl azide is significantly ($p<0.05$, $n=50$) higher compared to the controls. Furthermore, the difference between the bead sample exposed to unmodified cellopentaose and only PBS is insignificant ($p>0.05$, $n=50$). These results indicate a successful functionalization of beads with azido-labeled carbohydrates and the ability of type B CBMs to bind to the click-chemistry immobilized carbohydrates displayed on PS beads.

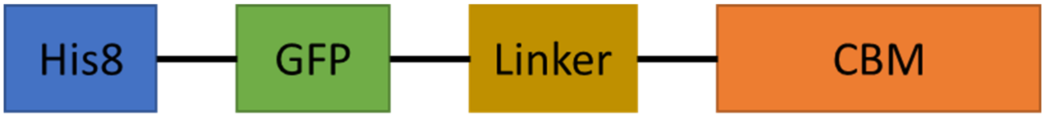


*Supplemental Figure S1: Domain overview of CBM fusion constructs. Both CBMs are fused to His$_8$-GFP via a flexible linker region. The individual sequences are found in Supplementary Table 1.*

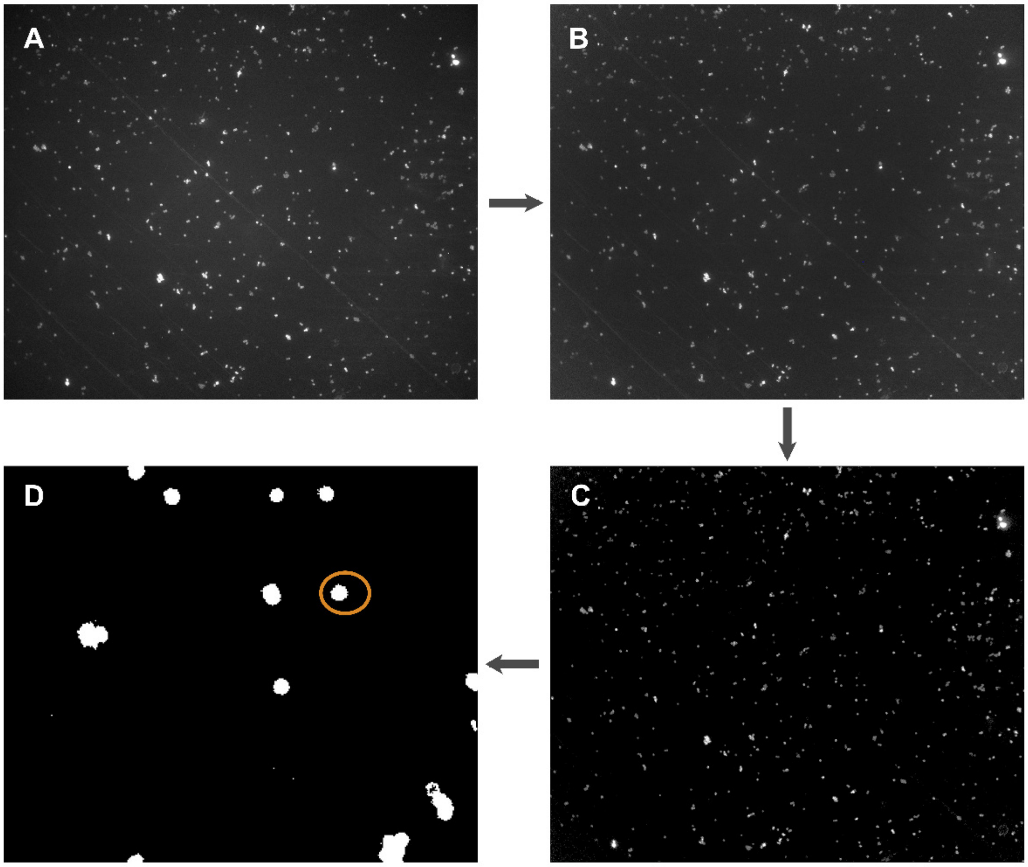


*Supplemental Figure S2: Single bead analysis flow chart. A) Raw image file. B) Image after background and flatfield correction. C) Image after thresholding and background removal. D) Only single, isolated beads such as the one circled in orange are analyzed.*

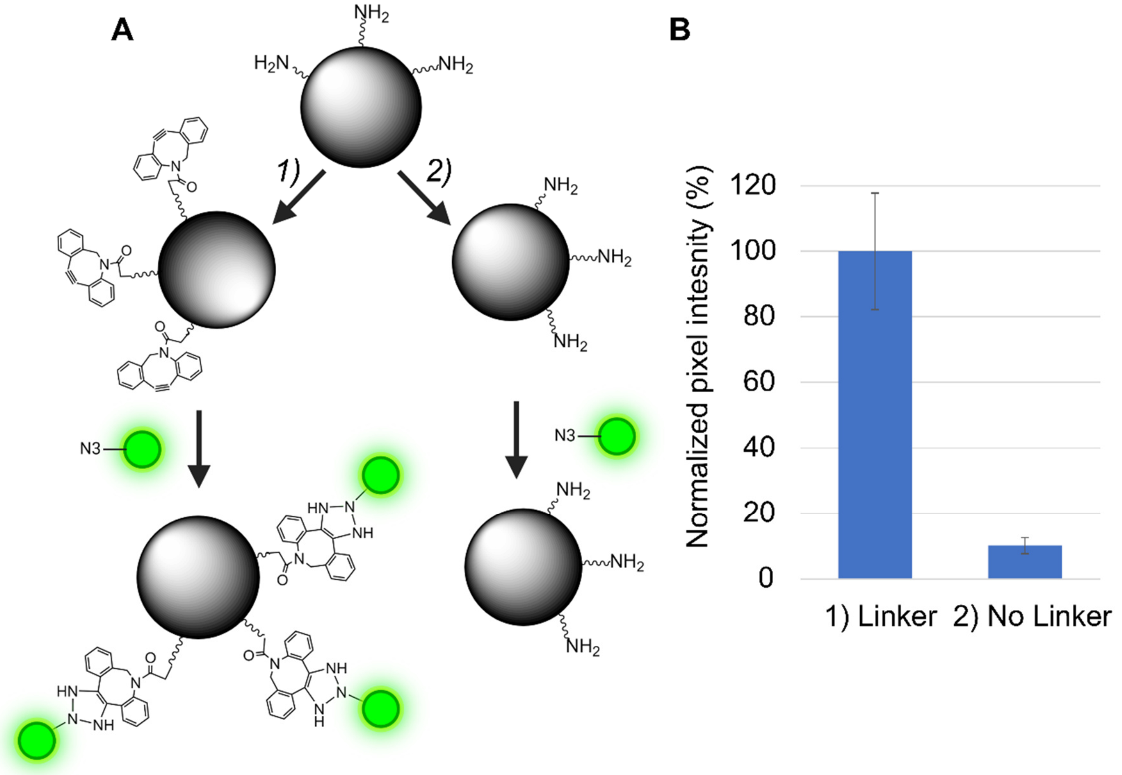


*Supplemental Figure S3: Verification of DBCO functionalization of aminated PS beads. A) Schematic overview of the experiment where in route 1) beads are functionalized with NHS-PEG(4)-DBCO and in route 2) beads are just incubated in PBS. B) Results of single bead fluorescence analysis. The fluorescence intensity is significantly ($p<0.05$, $n=50$) higher for DBCO functionalized beads. The pixel intensity is normalized to the intensity value of the linker-containing sample. Error bars represent the standard deviation of 50 analyzed beads.*

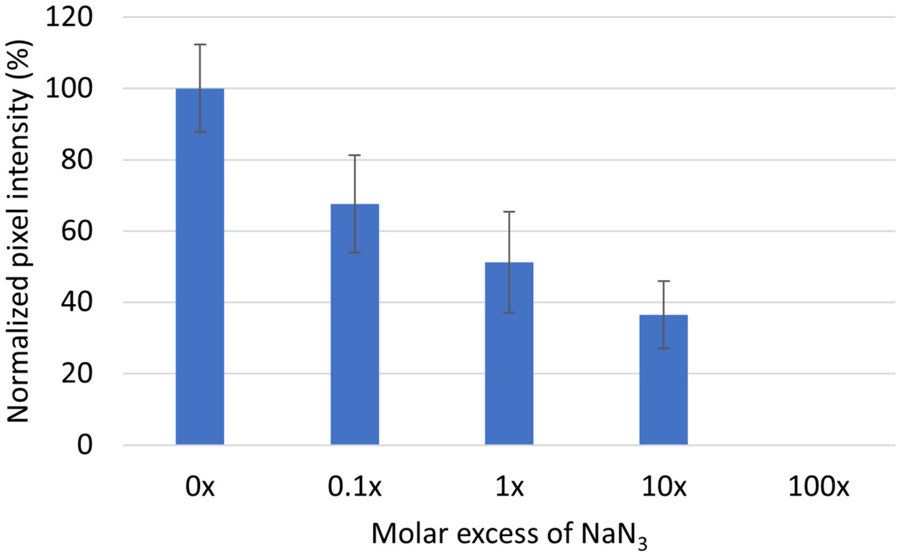


*Supplemental Figure S4: Influence of free azides on the click reaction between DBCO-modified beads and $N_3$-488. Even a small molar excess of free azides significantly ($p<0.05$, $n=50$) reduces the fluorescence signal, indicating a strong interference of free azides. The pixel intensity is normalized to the intensity value of the 0x molar excess sample. No fluorescence signal for the 100x sample was detected at the 300 ms exposure time. Error bars represent the standard deviation of 50 analyzed beads.*

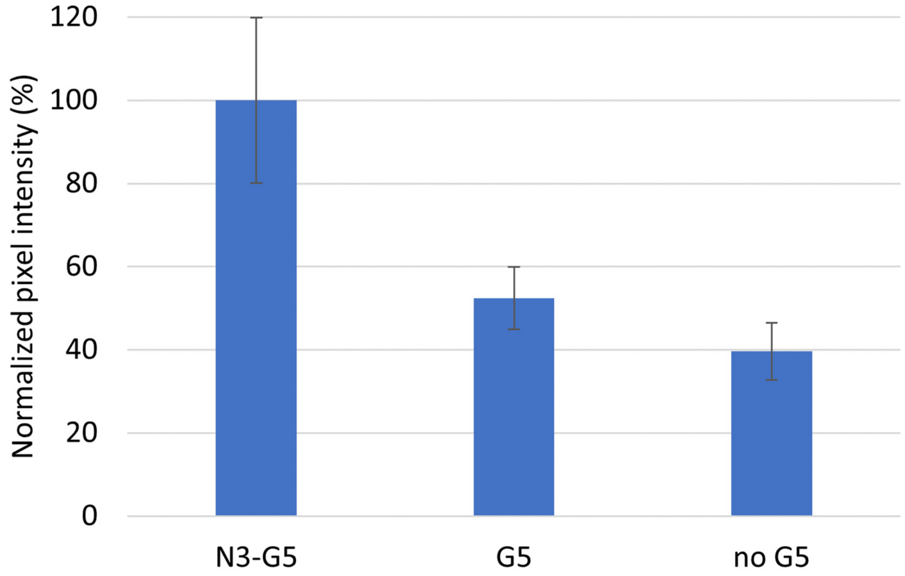


*Supplemental Figure S5: Type-B CBMs specifically bind to PS beads covalently functionalized with oligosaccharides. The obtained fluorescence intensity of carbohydrate modified beads is significantly ($p<0.05$, $n=50$) higher compared to the control experiments with unmodified cellopentaose or in the absence of any carbohydrate. The pixel intensity is normalized to the intensity value of the N3-G5 sample. Error bars represent the standard deviation of 50 analyzed beads.*

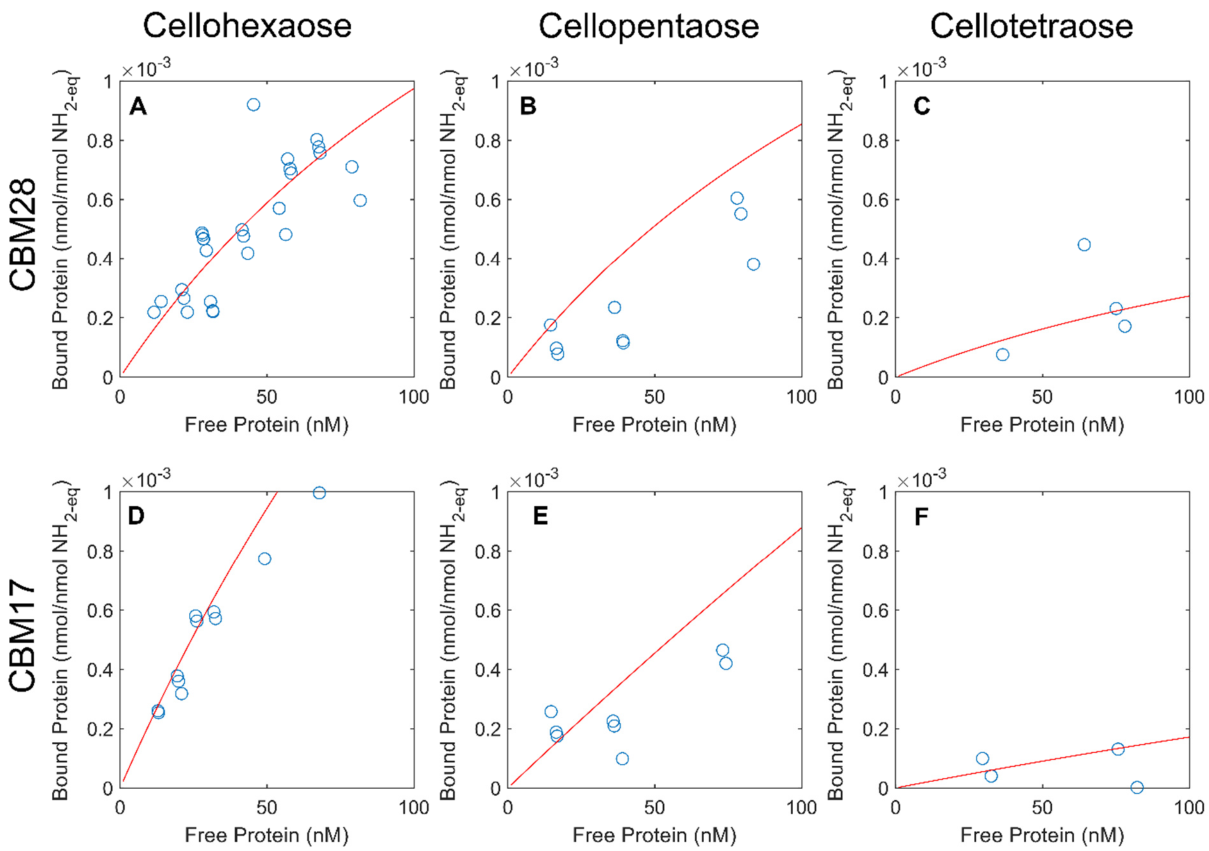


*Supplemental Figure S6: Binding data (blue circles) and fitted isotherms (red line) of CBM28 (top) and CBM17 (bottom) for cello-oligosaccharide-modified beads in the range 0-100 nM. Panel A-C) binding data for CBM28 on cellohexaose, cellopentaose and cellotetraose-modified beads respectively. Panel D-F) binding data for CBM17 on cellohexaose, cellopentaose and cellotetraose-modified beads respectively.*

*Supplemental Table S1: DNA sequences of linker domain, CBM17, and CBM28.*

| Domain | DNA sequence | Organism Source |
|---|---|---|
| Linker | GGTTTAAACGCGACTCCCACTAAAGGTGCCACTCCTACCAATA<br>CGGCGACTCCGACTAAGTCGGCAACGGCAACGCCCACTCGCC<br>CCAGCGTACCGACCAATACTCCGACTAATACCCCGGCGAACAC<br>CCTTAAG | N/A |
| CBM17 | AGCCAACCGACCGCCCCGAAAGATTTTTCCTCAGGTTTCTGGG<br>ACTTTAACGATGGCACGACCCAGGGTTTCGGCGTGAATCCGGA<br>CTCGCCGATTACGGCAATCAACGTTGAAAATGCTAACAATGCG<br>CTGAAAATTAGCAACCTGAACAGCAAAGGTAGTAACGATCTGTC<br>CGAAGGCAATTTTTGGGCCAACGTCCGCATCTCAGCAGACATT<br>TGGGGTCAATCGATCAATATTTATGGCGATACCAAACTGACGAT<br>GGACGTGATCGCTCCGACCCCGGTTAACGTCAGCATTGCGGC<br>CATCCCGCAGTCTAGTACGCATGGTTGGGGCAATCCGACCCGT<br>GCAATTCGCGTGTGGACGAACAATTTCGTTGCTCAAACCGATG<br>GTACGTATAAAGCGACCCTGACGATCTCCACCAACGACTCACC<br>GAATTTTAACACCATTGCCACCGATGCAGCCGACTCGGTCGTT<br>ACCAATATGATCCTGTTCGTGGGCTCCAACAGCGATAATATTAG<br>CCTGGACAACATCAAATTTACCAAA | *Clostridium cellulovarans* |
| CBM28 | ACCCAGAGCGGCACCGAAGTGGAAATTCCGGTGGTGCATGAT<br>CCGAAAGGCGAAGCGGTGCTGCCGAGCGTGTTTGAAGATGGC<br>ACCCGTCAGGGCTGGGATTGGGCGGGCGAAAGCGGCGTGAAA<br>ACCGCGCTGACCATTGAAGAAGCGAACGGCAGCAACGCGCTG<br>AGCTGGGAATTTGGCTATCCGGAAGTGAAACCGAGCGATAACT<br>GGGCGACCGCGCCGCGTCTGGATTTTTGGAAAAGCGATCTGG<br>TGCGTGGCGAAAACGATTATGTGACCTTTGATTTTTACCTGGAC<br>CCAGTGCGTGCGACCGAAGGCGCGATGAACATTAACCTGGTGT<br>TTCAGCCGCCGACCAACGGCTATTGGGTGCAGGCGCCGAAAA<br>CCTATACCATTAACTTTGATGAACTGGAAGAACCGAACCAGGTG<br>AACGGCCTGTATCATTATGAAGTGAAAATTAACGTGCGTGATAT<br>TACCAACATTCAGGATGATACCCTGCTGCGTAACATGATGATTA<br>TTTTTGCGGATGTGGAAAGCGATTTTGCGGGCCGTGTGTTTGT<br>GGATAACGTGCGTTTTGAAGGCGCG | *Alkalihalobacillus akabai (Bacillus sp. 1139)* |

*Supplemental Table S2: p-values for fit parameters corresponding to main text Table 2. The p-values for cellotetraose are > 0.05 indicating a weak fit of the model.*

| | **Cellohexaose-beads** | | **Cellopentaose-beads** | | **Cellotetraose-beads** | |
|---|---|---|---|---|---|---|
| | $K_d$ | $n_{max}$ | $K_d$ | $n_{max}$ | $K_d$ | $n_{max}$ |
| **CBM28** | 1.29E-07 | 1.33E-16 | 0.05027 | 6.18E-05 | 0.57787 | 0.14801 |
| **CBM17** | 4.11E-04 | 4.69E-09 | 0.0336 | 0.00134 | 0.56302 | 0.33078 |
